# DNA Compression with Genomic Language Models: Tokenization, Benchmarking, and an Information-Content Map

**DOI:** 10.64898/2026.06.10.731316

**Authors:** Vojtech Macala, Petr Simecek

## Abstract

Lossless compression and probabilistic sequence modeling are two faces of the same coin: a model that assigns high probability to a sequence can encode it in few bits via arithmetic coding. We exploit this duality to evaluate genomic language models as compressors of DNA, using compression primarily as an objective probe of generative sequence modeling rather than as a deployable storage system. We release DNAGPT2, a family of ten GPT-2-small models pretrained for one epoch on a single A40 using the DNABERT2 multi-species corpus that differ only in byte-pair encoding vocabulary size. Coupled with arithmetic coding, the best model reaches 1.47 bits per base (bpb) on the T2T human genome, fourth in the Cobilab compression benchmark and ahead of every general-purpose compressor. Our results suggest that NLP-style tokenization choices may be suboptimal for DNA: a 32-token BPE vocabulary compresses better than larger vocabularies. We also find that, in this benchmark, published long-context genomic LMs underperform a much shorter-context BPE GPT-2; we discuss in Section 5 that this is not a controlled context-length ablation, since the compared models also differ in architecture, training data, parameter count, and tokenization. Finally, we compute a per-nucleotide information-content map of the human genome and show that exons, introns, intergenic regions, and Alu repeats have statistically distinct information profiles.

## 1. Introduction

Genomic sequence data is being produced faster than storage capacity grows. Compression therefore matters in practice, but it is also a clean quantitative probe of generative models. A probabilistic model with cross-entropy *H* can in principle compress its input down to *H* bits per symbol via arithmetic coding; the compression ratio is the model’s predictive quality made tangible (Delétang et al., 2024; Witten et al., 1987).

DNA is a particularly interesting target for this view. Its alphabet has only four symbols, coding regions occupy less than 3% of the human genome, and much of the remaining sequence is shaped by repeats, transposons, and local composition biases. A strong generative model of DNA should therefore be both a compressor and a source of biologically meaningful “surprise” tracks.

Recent genomic language models such as DNABERT2, HyenaDNA, megaDNA, and CD-GPT have demonstrated strong performance on sequence modeling and downstream genomic benchmarks (Benegas et al., 2025; Zhou et al., 2024; Nguyen et al., 2023; Shao & Yan, 2024; Zhu et al., 2024). Their predictive sharpness on raw DNA, however, has not been systematically measured against the mature literature of specialized DNA compressors. This paper uses lossless compression as a model-evaluation lens and then turns per-token surprise into a biological track for genome exploration.

To study this systematically, we train and release DNAGPT2, a family of ten GPT-2-small genomic language models that are identical except for BPE vocabulary size (16–8192 tokens). We then benchmark DNAGPT2, CD-GPT, HyenaDNA, and megaDNA as probability models for arithmetic coding, comparing them against strong DNA compressors and general-purpose baselines on three genomes. This benchmark exposes two notable patterns: small BPE vocabularies can compress DNA better than larger ones, and published long-context genomic LMs do not automatically yield lower next-token entropy. Finally, we use

DNAGPT2-derived surprise to build a genome-wide information-content map of GRCh38, where exons, introns, intergenic regions, and Alu repeats separate into distinct information profiles; the resulting tracks are released as an interactive genome-browser resource.

We stress from the outset that this is an evaluation and analysis study, not a practical compressor proposal. The compressed sizes we report measure the log-likelihood that a generative model assigns to a full genome, but they exclude model weights and assume bit-exact decoder synchronization. We maintain this distinction throughout: compression serves as a lens for what genomic LMs learn, and per-token surprise serves as a candidate biological signal.

## 2. Background and Related Work

Predictive coding with LMs. Arithmetic coding takes a stream of conditional distributions *p*(*x*_*t*_ | *x*_*<t*_) and emits a bitstream of length close to ∑_*t*_ − log_2_ *p*(*x*_*t*_ |*x*_*<t*_) (Witten et al., 1987). Any autoregressive language model is therefore directly usable as a compressor, given a synchronized decoder. Deletang et al. showed this relationship across modalities (Delétang et al., 2024). NNCP trains a transformer online on the data being compressed and leads several general-purpose benchmarks (Bellard, 2021). Our setting is offline: a single pretrained model serves as a fixed probability source.

This offline setting is closer to standard foundation-model evaluation than to classic adaptive compression. The model is trained once on a large corpus and then evaluated on held-out or partially held-out genomes. In exchange for worse practical compression overhead, we get a clean quantity: the number of bits required to encode an entire genome under the model’s conditional distribution. This quantity is comparable across model families and does not depend on task-specific labels, making it attractive for genomic generative models whose downstream benchmarks are often heteroge-neous.

DNA compressors. DNA compressors range from general-purpose tools (gzip, bzip2, lzma, paq8l) to specialized reference-based and reference-free algorithms (Gilmary & Sharma, 2023). State-of-the-art reference-free tools combine finite-context models, repeat models, and shallow neural mixers: GeCo3 (Pratas et al., 2020), JARVIS3 (Sousa et al., 2024), and MFCompress (Pinho & Pratas, 2014). Neural DNA-compression work such as DeepDNA has also explored next-symbol prediction with neural networks, but mostly on short or organelle-scale genomes (Silva et al., 2020). On the standardized T2T human-genome benchmark maintained by Cobilab, JARVIS3 currently leads at 1.384 bpb (Computational Biology Lab, University of Aveiro, 2024). Our work explores a different point in this design space: a much larger autoregressive neural model with no hand-engineered repeat module.

The contrast with JARVIS3 is especially informative. JARVIS3 combines local Markov-style context models with explicit repeat models and a neural mixing component. That design encodes two empirical facts about genomes: local composition is highly informative, and long exact or approximate repeats matter a lot. Decoder-only genomic LMs instead rely on their learned hidden state to represent both effects. If such models underperform specialized compressors, the gap helps identify what architectural bias is missing.

Tokenization of DNA. How DNA is tokenized before being fed to a language model is a design choice with significant consequences. Three broad families exist: (i) nucleotide-level tokenization maps each base to one of four token ids—used by HyenaDNA and megaDNA—preserving SNP resolution but producing very long sequences; (ii) *k*-mer tokenization groups *k* consecutive nucleotides, either overlapping (DNABERT, *k* = 6) (Ji et al., 2021) or non-overlapping (Nucleotide Transformer), reducing sequence length at the cost of shift sensitivity; (iii) byte-pair encoding (BPE) learns a data-driven vocabulary of variable-length subwords (Kudo & Richardson, 2018)—used by CD-GPT and by our DNAGPT2 family. In NLP, BPE vocabularies of 50k–128k tokens are standard (Radford et al., 2019; Grattafiori et al., 2024); whether such sizes are appropriate for the four-letter DNA alphabet is one of the questions we address.

Genomic language models. DNABERT2 introduced a multi-species pretraining corpus of 135 genomes, which we reuse for DNAGPT2 (Zhou et al., 2024). Among decoder-only models suitable for compression, HyenaDNA replaces attention with the Hyena long-convolution operator and single-nucleotide tokens, en-abling contexts up to 1M bases while remaining compact (Nguyen et al., 2023). megaDNA adopts the MEGABYTE multiscale architecture (Yu et al., 2023) with three hierarchical layers—local, middle, and global—trained on bacteriophage genomes (Shao & Yan, 2024). CD-GPT is a multi-omic GPT trained on DNA, RNA, and proteins with a shared BPE vocabulary (Zhu et al., 2024). Several recent genomic LMs are not included in our head-to-head benchmark: masked models such as the Nucleotide Transformer (Dalla-Torre et al., 2023) and bidirectional models such as Caduceus (Schiff et al., 2024) do not directly yield causal next-token distributions for arithmetic coding without additional adaptation. We therefore restrict the benchmark to decoder-only models that natively expose conditional next-token probabilities.

## 3. Methods

### 3.1. DNAGPT2

We train ten autoregressive transformers sharing the GPT-2-small architecture: 12 layers, 12 attention heads, hidden size 768, context length 1024, and approximately 86M parameters (Radford et al., 2019). The models differ only in BPE vocabulary size *V* ∈{16, 32, 64, 128, 256, 512, 1024, 2048, 4096, 8192}.

We use the DNABERT2 multi-species corpus (Zhou et al., 2024), filtered to A/C/G/T, dropping ambiguous N positions. After filtering, the corpus contains approximately 32.5B nucleotides spanning 135 genomes. Each SentencePiece BPE tokenizer is trained on one sixth of the corpus (Kudo & Richardson, 2018). All models are trained for one epoch (≈10.5B tokens for the *V* = 128 tokenizer) with AdamW, cosine learning-rate decay, total batch size 2^19^ tokens, sequence length 1024, bf16 mixed precision, and a single NVIDIA A40 GPU.

Figure 3 shows the training and validation curves for DNAGPT2_128, the model used for genome-wide information-content mapping. The curves are monotone and closely matched, suggesting that underfitting rather than overfitting is the dominant limitation. This matters for interpreting the benchmark: the study isolates tokenizer effects under a fixed training budget, but absolute bpb numbers would likely improve with longer training or larger models.

### 3.2 Compression Pipeline

The pipeline (Figure 1) tokenizes the input sequence, processes token windows of length 1024 with stride 512, converts next-token logits to probabilities and cumulative distribution functions (CDFs), quantizes the CDFs to integers with scale 2^15^, enforces a minimum interval of one, and arithmetic-codes each token. Decompression reverses the process using the same weights, tokenizer, and quantization scheme. Appendix Figure 9 shows a worked arithmetic-coding example.

**Figure 1.**
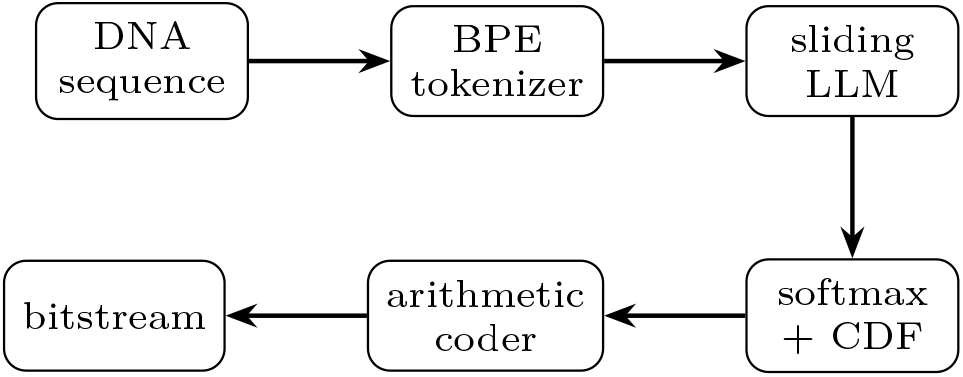
Compression pipeline. DNA is tokenized, processed by a causal genomic LM in sliding windows, converted to integer CDFs, and arithmetic-coded. Decoding uses the same model and quantization scheme.

The sliding-window stride controls a speed/accuracy trade-off. A stride equal to the context length would evaluate each token with little overlap between neigh-boring windows and therefore less left context at window boundaries. Smaller strides improve conditioning but require more forward passes. We use half-context stride as a practical compromise. CDF quantization is another important implementation detail: arithmetic coders operate over integer frequency intervals, so extremely small softmax probabilities can vanish after rounding. Enforcing a minimum interval of one prevents impossible symbols during decoding, at the cost of a small amount of probability-mass distortion.

### 3.3 Models and Datasets

We evaluate four genomic LMs: DNAGPT2_32 (BPE 32, 1024 context, 86M parameters), CD-GPT (BPE 64k, 1024 context, 1B parameters), HyenaDNA-medium (single-nucleotide, 160k context, 6.5M parameters), and megaDNA (96k context, 145M parameters). Classical baselines include JARVIS3, GeCo3, MFCompress, bzip2, and gzip; for human we also reproduce Cobilab benchmark entries.

These models span several axes that matter for compression (Table 1). DNAGPT2_32 and CD-GPT share a 1024-token context but differ in vocabulary size, parameter count, and training scope. HyenaDNA and megaDNA use single-nucleotide or byte-level representations and much longer contexts, but differ in architecture: HyenaDNA uses implicit long convolu-tions, while megaDNA uses a hierarchical multiscale transformer. The comparison is therefore deliberately broad rather than a controlled ablation; its purpose is to test whether published generative DNA models provide useful probability distributions for entropy coding.

**Table 1.**
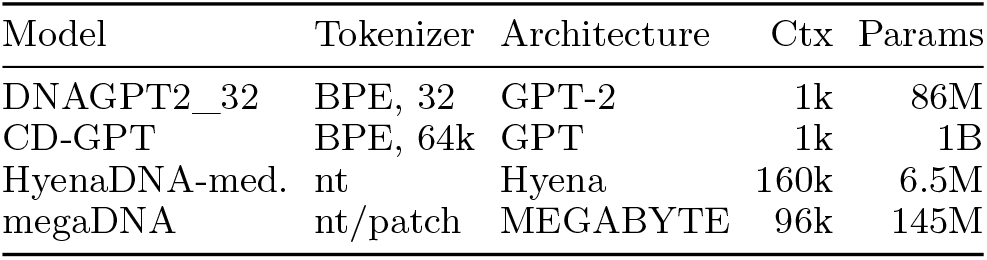
Genomic LMs evaluated as probability models for arithmetic coding.

Compression is measured on three references: Homo sapiens T2T-CHM13v2.0 (Nurk et al., 2022), Myxococcus llanfairpwllgwyngyllgogerychwyrndrobwllllantysiliogogogochensis^1^ (Chambers et al., 2020), and Arabidopsis thaliana chromosome 1 (The Arabidopsis Information Resource). Human sequence appears in the DNABERT2 pretraining corpus, so the T2T benchmark should be treated as partly in-distribution for DNAGPT2; M. llanfair… and A. thaliana check cross-species behavior.

The three datasets are intentionally diverse. The human genome gives direct comparability to the Cobilab leaderboard. The bacterial genome tests cross-domain behavior on a compact prokaryotic sequence. The plant chromosome tests a eukaryotic but non-animal genome with different repeat and GC-composition structure. The goal is not to exhaustively characterize generalization, but to ensure that the main findings are not artifacts of a single human benchmark.

### 3.4. Information Content

For each token *x*_*t*_ of length *𝓁*_*t*_, we define information content

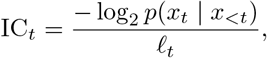

in bits per base. We compute IC over GRCh38.p14 using DNAGPT2_128. Its compression cost over the *V* = 32 optimum is negligible (Table 2), while its larger vocabulary reduces the per-genome token count, lowering the cost of the genome-wide IC pass. Values are stored as bedGraph and bigWig tracks at 25, 250, and 2500 bp resolutions (UCSC Genome Browser Group, a;b). We intersect tracks with GENCODE v48 exons, introns, intergenic regions, and RepeatMasker Alu repeat coordinates (Frankish et al., 2025). Group differences are tested with Kruskal-Wallis and pairwise Dunn tests with Bonferroni correction on *N* = 5 × 10^6^ sampled tokens.

**Table 2.**
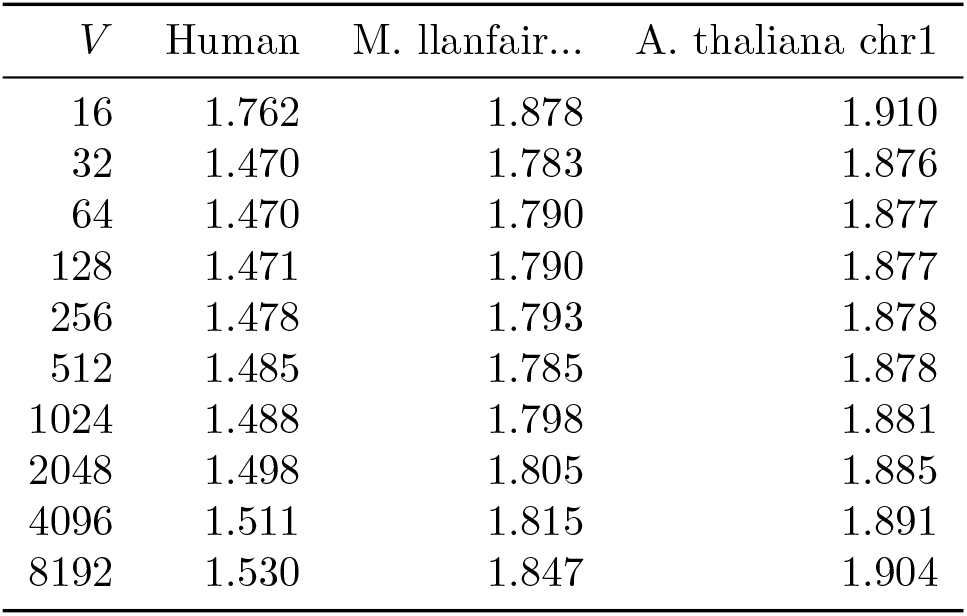
DNAGPT2 compression in bits per base across vocabulary sizes and genomes. Bold marks the best vocabulary per genome.

## 4. Results

### 4.1. Tokenization: Smaller is Better

Before neural modeling, BPE tokenization is itself a pre-compression step. If *n* nucleotides become *T* tokens under vocabulary size *V*, fixed-width token coding costs

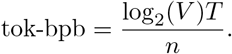

The left panel of Figure 2 shows a U-shape: the tokenization-only optimum is at *V* = 128, because longer vocabularies yield diminishing gains in average token length while their per-token cost grows linearly.

**Figure 2.**
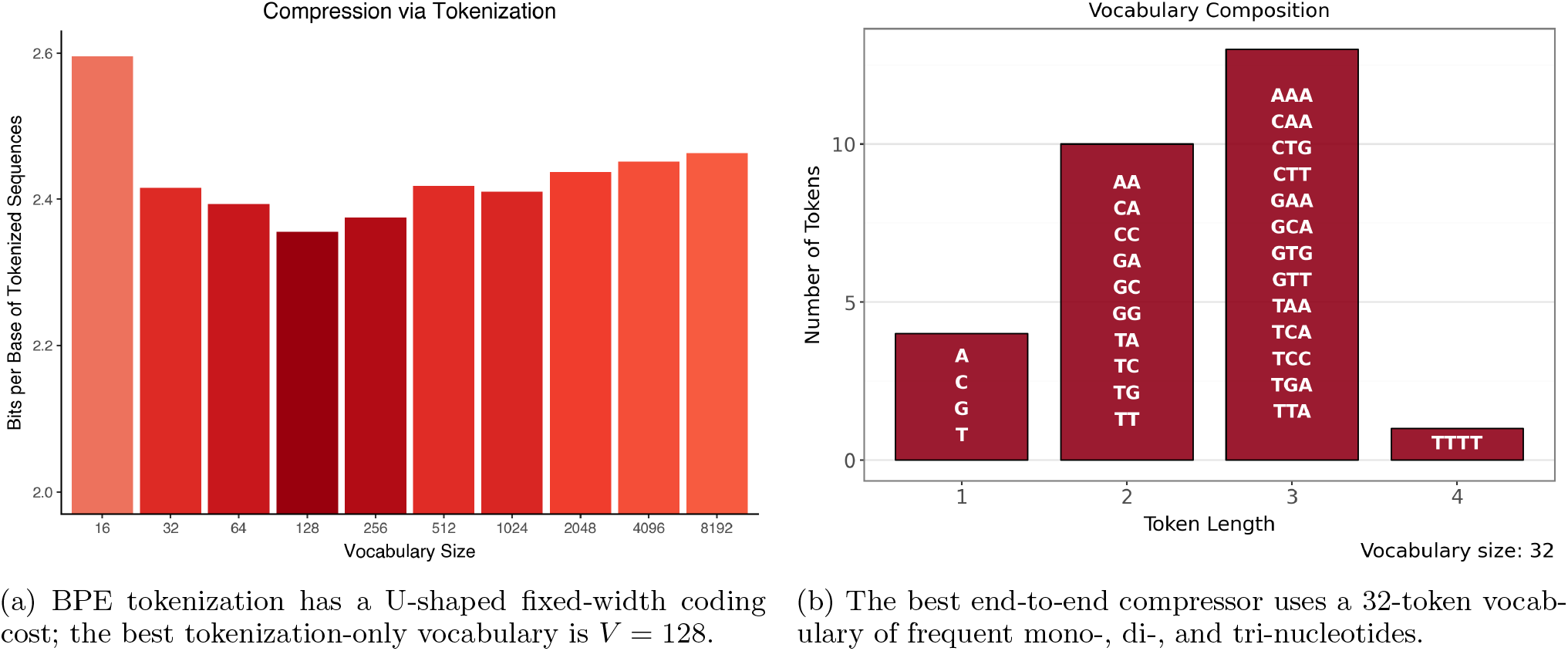
Effect of BPE vocabulary size on genomic compression. Left: fixed-width coding cost of tokenized sequences, averaged over the three benchmark genomes, is minimized at *V* = 128. Right: the vocabulary that gives the best end-to-end DNAGPT2 compression is much smaller (*V* = 32) and consists almost entirely of mono-, di-, and tri-nucleotides.

**Figure 3.**
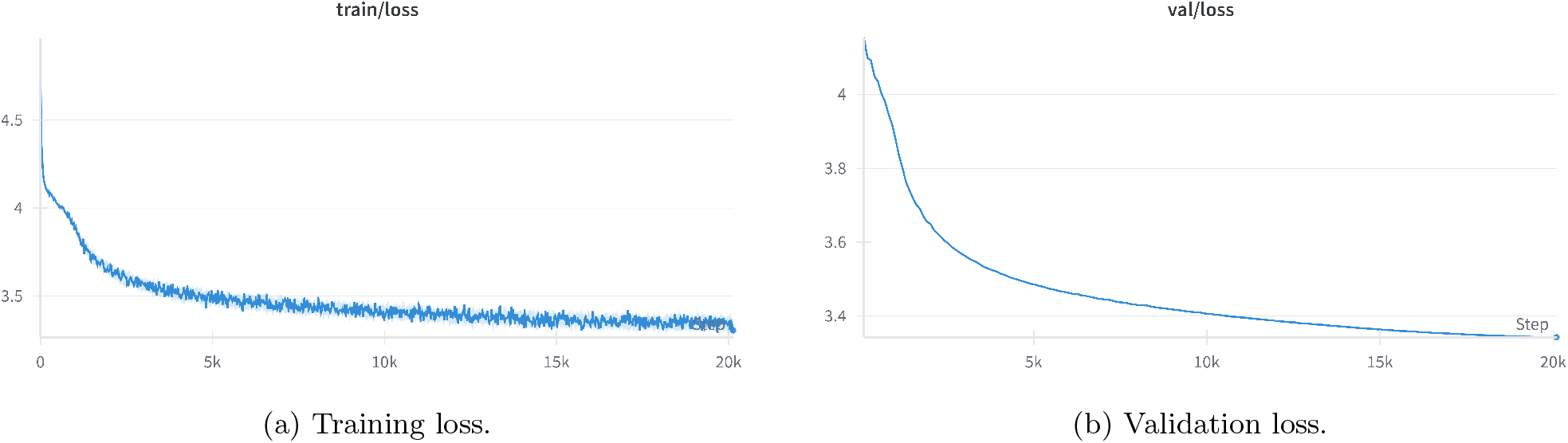
DNAGPT2_128 loss curves over one epoch (*∼*20k optimizer steps). Loss is still decreasing at the end of training, so the reported compression numbers should be interpreted as a compute-constrained first pass rather than a fully saturated model family.

The diagnostics in Figure 4 explain the U-shape. Doubling *V* always costs one additional bit per token under fixed-width coding, but the corresponding gain in nucleotides per token becomes progressively smaller. At large *V*, much of the vocabulary is occupied by rare long motifs. These motifs shorten the sequence only modestly and are difficult for a fixed-capacity model to predict because the training signal for each rare token is sparse.

**Figure 4.**
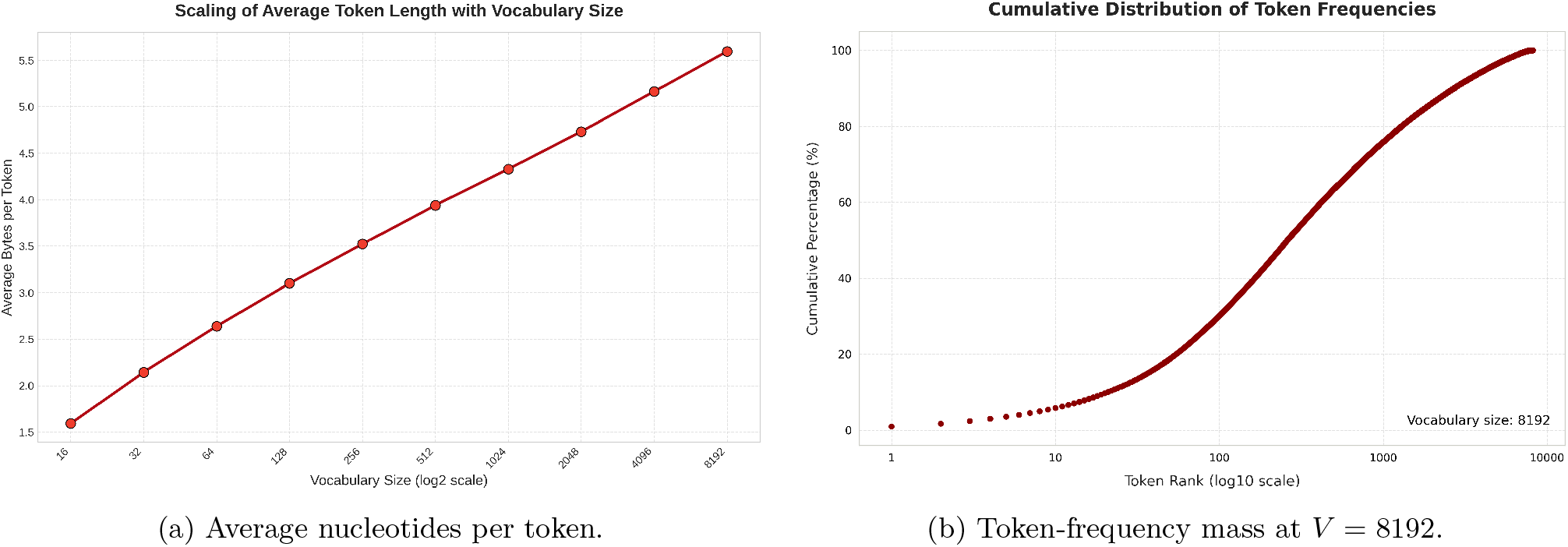
Tokenizer scaling diagnostics. Left: average token length grows sub-linearly with vocabulary size, so each vocabulary doubling costs one extra bit per token but yields progressively smaller sequence-length savings. Right: at *V* = 8192, the token distribution has a long tail, with the top *∼*1000 tokens accounting for most observed tokens.

When the full compression pipeline is applied (LLM inference followed by arithmetic coding), the optimum shifts to *V* = 32 (Table 2). This vocabulary wins on all three genomes.

The winning tokenizer is unexpectedly simple (Figure 2, right): it is essentially a mixed 1–3-mer alphabet plus the homopolymer TTTT. Its tokens are frequent and well-supported, making prediction easier for a fixed-capacity model. Larger vocabularies introduce longer, rarer tokens that are under-trained under our one-epoch budget.

This result separates two effects that are easy to conflate. A tokenizer alone can shorten a sequence by packing frequent motifs into longer symbols; by that metric, *V* = 128 is best. A compressor, however, pays not only for sequence length but also for the model’s uncertainty over the next token. Larger BPE vocab-ularies reduce token count but make the prediction problem harder because probability mass is spread over more rare tokens. For the fixed DNAGPT2 architecture and one-epoch training budget, the predictive gain from the smaller vocabulary dominates the pre-compression gain from the larger vocabulary.

### 4.2. Compression Benchmark

The following comparison reports sequence bitstreams only; it excludes model weights, tokenizer files, and the burden of bit-exact decompression. It should therefore be read primarily as model evaluation, not as a claim that LLMs are deployable drop-in DNA compressors.

On the partly in-distribution human benchmark, DNAGPT2_32 reaches 1.470 bpb (Table 3), fourth in the Cobilab ranking and only about 6% behind JARVIS3. It also slightly outperforms NNCP, a transformer-based compressor that adapts online to the input. The two non-human genomes in Table 4 are therefore important checks: DNAGPT2_32 remains the best LLM on A. thaliana, while megaDNA is narrowly best among LLMs on the bacterium.

**Table 3.** T2T-CHM13v2.0 human genome compression benchmark. Bold rows are genomic LMs evaluated in this work; other rows are best-of-runs reported by Cobilab.^1 2^.

| Program | bpb | Time (min) | Factor | Approach |
| --- | --- | --- | --- | --- |
| JARVIS3 | 1.384 | 641 | 31% | specialized + ML |
| JARVIS2 | 1.395 | 381 | 30% | specialized stat. |
| GeCo3 | 1.425 | 690 | 29% | specialized + ML |
| DNAGPT2_32 | 1.470 | 728 | 26% | genomic LLM |
| CD-GPT | 1.511 | 8665 | 24% | genomic LLM |
| NNCP | 1.514 | 17465 | 24% | online transformer |
| MFCompress | 1.560 | 22 | 22% | specialized stat. |
| GeCo2 | 1.560 | 48 | 22% | specialized stat. |
| JARVIS | 1.564 | 171 | 22% | specialized stat. |
| paq8l | 1.571 | 4588 | 21% | general-purpose |
| bsc-m03 | 1.577 | 39 | 21% | general-purpose |
| NAF-22 | 1.640 | 43 | 18% | specialized stat. |
| lzma -9 | 1.658 | 84 | 17% | dictionary |
| HyenaDNA | 1.790 | 2286 | 10% | genomic LLM |
| megaDNA | 1.848 | 8252 | 7% | genomic LLM |
| bzip2 -9 | 1.932 | 5 | 3% | dictionary |
| gzip -9 | 2.022 | 56 | -1% | dictionary |

**Table 4.** Cross-genome compression in bpb. Bold marks the best overall; italic marks the best LLM.

| Method | Human | M. llanfair... | A. thaliana |
| --- | --- | --- | --- |
| DNAGPT2_32 | 1.470 | 1.783 | 1.876 |
| CD-GPT | 1.511 | 1.769 | 1.886 |
| HyenaDNA | 1.790 | 2.219 | 1.962 |
| megaDNA | 1.848 | 1.755 | 1.923 |
| JARVIS3 | 1.384 | 1.713 | 1.702 |
| GeCo3 | 1.425 | 1.741 | 1.728 |
| MFCompress | 1.560 | 1.826 | 1.819 |
| bzip2 -9 | 1.932 | 2.077 | 2.112 |
| gzip -9 | 2.022 | 2.142 | 2.161 |

DNAGPT2_32 also outperforms the longer-context HyenaDNA and megaDNA on all three genomes. This is not a controlled context-length ablation: these models differ in architecture, tokenization, data, objective, and implementation. The narrower conclusion is that published long-context genomic LMs do not automatically translate their larger context windows into lower next-token entropy. LLM compression is also slow: DNAGPT2 takes 728 minutes on human, compared with 22 minutes for MFCompress.

The benchmark therefore has two complementary messages. On the positive side, a relatively small GPT-style model trained for a single epoch produces probabilities strong enough to compete with mature specialized compressors. On the negative side, the remaining gap to JARVIS3 is meaningful, and the runtime gap is large. We view this as evidence that genomic LMs are already useful as probabilistic models, but that practical DNA compression will likely require hybridization with classical compressor mechanisms, especially explicit copy/repeat handling.

### 4.3. Information Content of the Human Genome

The compression results above measure aggregate model quality; the information-content analysis below asks a finer question: where in the genome does the model find sequence predictable or surprising? A key distinction from standard conservation tracks is that IC is model-relative and local. A high PhyloP score means a region evolved more slowly than expected across species; a high IC value means that, given the preceding sequence, DNAGPT2 assigns low probability to the observed token. These are related but not equivalent signals: repetitive elements can be low IC even when not conserved, while exons can be both conserved and locally difficult to predict.

DNAGPT2-derived information content separates annotated genomic features (Table 5, Figure 5). Exons have the highest median IC (1.875 bpb), followed by introns (1.814), intergenic regions (1.767), and Alu repeats (0.631). A Kruskal-Wallis test rejects equal medians and all pairwise Dunn tests remain significant after Bonferroni correction (*p <* 10^*−*3^).

**Table 5.** Information content by genomic feature on GRCh38 (bpb).

| Feature | Mean | Median | Std | 25% | 75% |
| --- | --- | --- | --- | --- | --- |
| Exon | 1.766 | 1.875 | 0.381 | 1.767 | 1.952 |
| Intron | 1.541 | 1.814 | 0.563 | 1.279 | 1.915 |
| Intergenic | 1.417 | 1.767 | 0.645 | 0.899 | 1.902 |
| Alu repeats | 0.725 | 0.631 | 0.439 | 0.396 | 0.962 |

**Figure 5.**
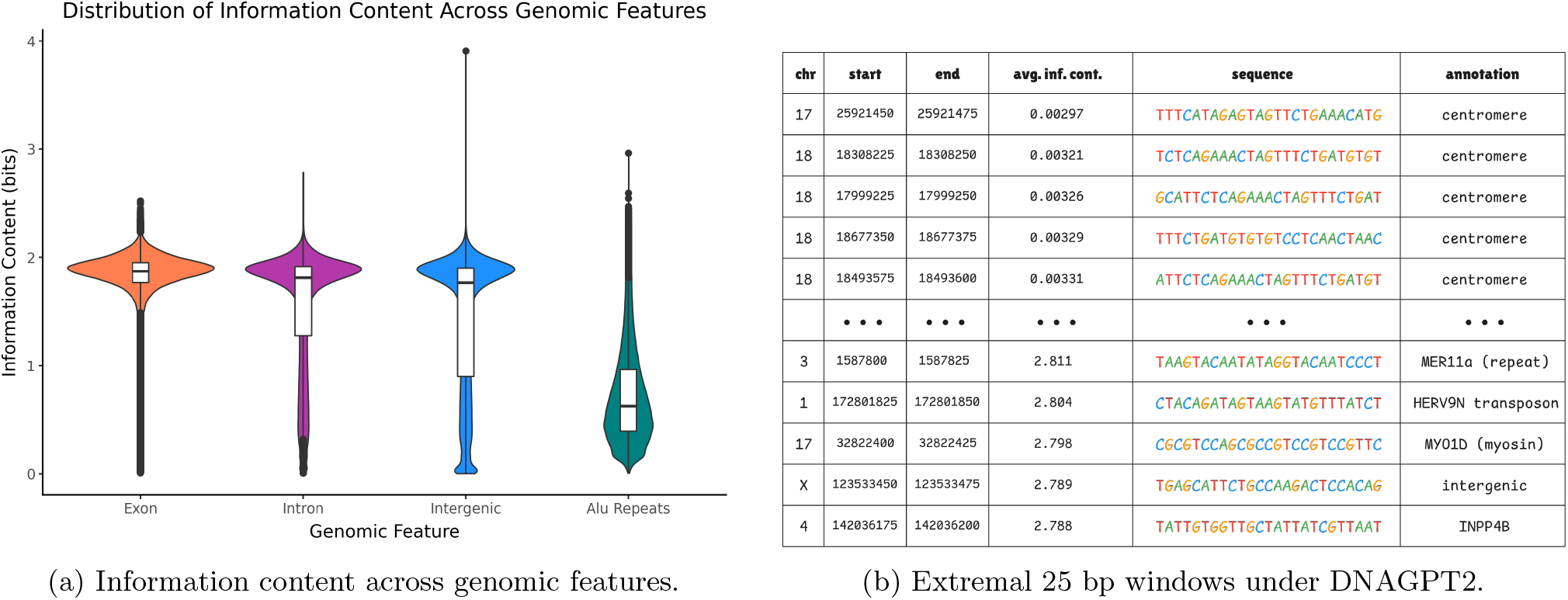
Information-content profiles of the human genome under DNAGPT2_128. Left: Alu repeats are substantially more predictable than exons, introns, or intergenic regions. Right: the lowest-information 25 bp bins concentrate in centromeric tandem repeats, while high-information bins occur in introns, transposons, and intergenic sequence.

The extremes make the model’s behavior concrete (Figure 5, right). The lowest-IC bins all lie in centromeric tandem repeats on chromosomes 17 and 18, which are nearly free under the model. Highest-IC bins occur in introns of MYO1D and INPP4B, a MER11a repeat, an HERV9N transposon, and an intergenic region on chrX.

These extremes are useful sanity checks. Centromeres were among the most challenging regions for complete human-genome assembly, precisely because they contain long, structured tandem repeats. DNAGPT2 assigns them very low information content because the next token is almost determined by local context. The high-IC examples are more heterogeneous: they include unique intronic sequence, repeat-derived elements, and intergenic sequence. This mixture is expected, because high IC is not a direct functional label; it marks regions that the model finds locally surprising.

The full track set is deployed in a JBrowse2 application (Figure 6), allowing users to explore how model surprise aligns with genes, repeats, and conservation tracks. An accompanying analysis interface (Appendix Figure 8) lets users query specific genes or repeat families and compare their IC distributions. Once the IC tracks are computed, arbitrary annotation sets can be overlaid without additional model inference, making the approach applicable to new genome assemblies or model variants with minimal effort.

**Figure 6.**
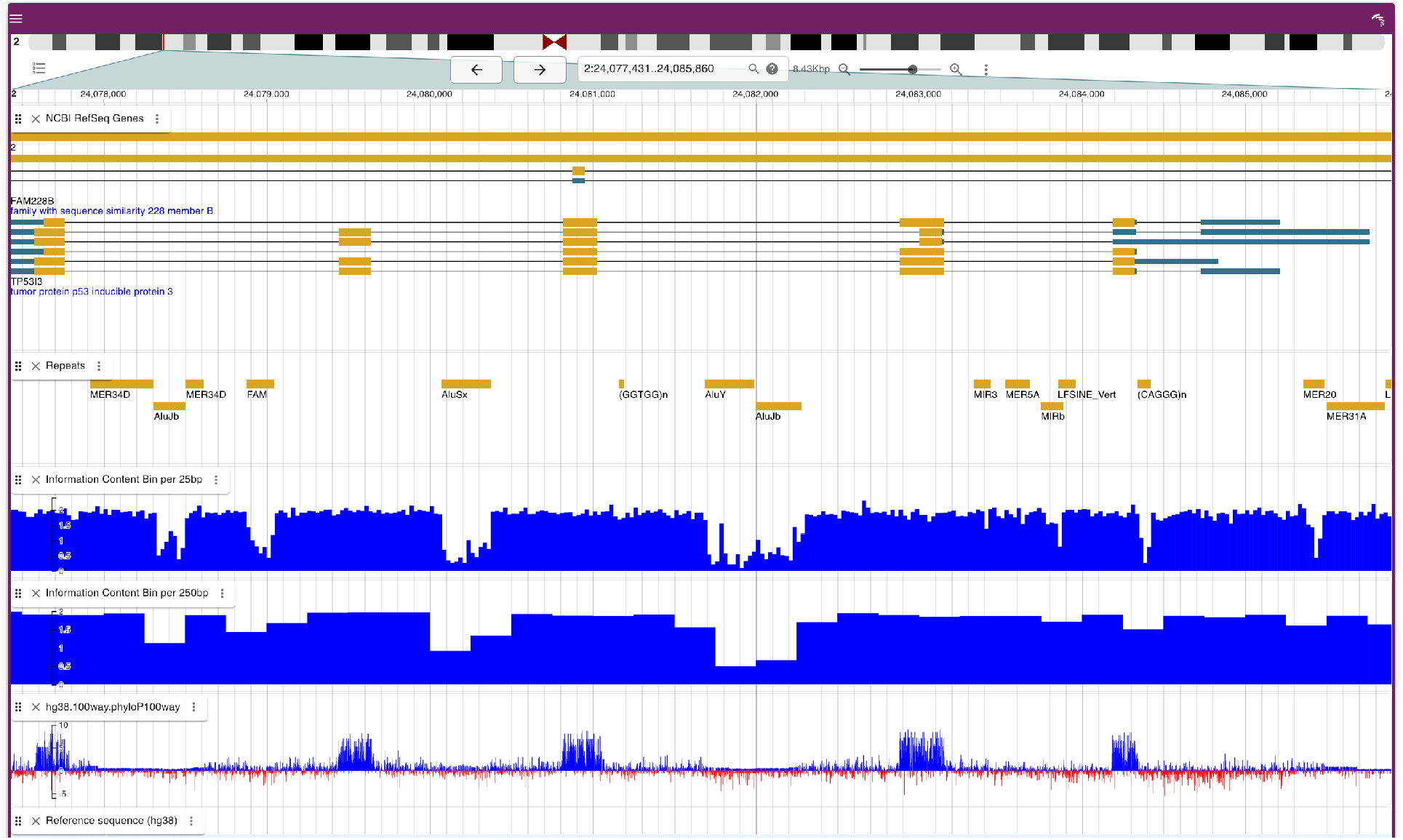
JBrowse2 view of DNAGPT2-derived information content on chromosome 2, aligned with RefSeq genes (Goldfarb et al., 2025), RepeatMasker elements, 25/250 bp information-content tracks, and PhyloP100way conservation (Pollard et al., 2010).

## 5. Discussion

Why might long context not translate into better compression here? The gap between DNAGPT2_32 and the longer-context HyenaDNA and megaDNA is not what a simple “more context is always better” story would predict. With the caveat that this is not a controlled ablation, two hypotheses are plausible. First, much next-nucleotide entropy may be dominated by short-range structure: di- and tri-nucleotide composition, codon bias, and local repeats. Long-range dependencies certainly exist in genomes, but may contribute less to local predictive entropy than to function, a distinction also raised in recent analyses of sequence-based regulatory models (Karollus et al., 2023). Second, top specialized compressors such as JARVIS3 combine local finite-context models with explicit repeat models, whereas the long-context LMs tested here lack an explicit copy mechanism.

Why might a small vocabulary win? The DNA alphabet has only four symbols, and BPE merges quickly converge to high-frequency *k*-mers. A 32-token vocabulary captures mono-, di-, and tri-nucleotide structure while keeping every token frequent. Larger vocabularies introduce lower-frequency merges that are under-trained at this model scale and training budget. Our results suggest that NLP defaults of 50k–128k tokens should not be transplanted into genomic compression without justification.

Compression as a genomic probe. The information-content map is a direct readout of what the model has learned. That exons, introns, intergenic regions, and Alu repeats separate without supervised labels suggests that a genomic LM trained only on raw sequence recovers known structure. We view IC tracks as complementary to conservation scores in genome browsers.

This complementarity is the main biological argument for the paper. Conservation summarizes evolutionary constraint across species, while IC summarizes predictability under a learned sequence model. A repet-itive region can be easy for the model but not conserved; a constrained exon can be conserved and still locally surprising. In practice, the most useful view may be the joint one: regions with high conservation and high IC are likely to be constrained and sequence-specific, whereas regions with low IC often reflect repetitive or low-complexity structure.

What LMs add beyond classical compressors. The benchmark does not suggest replacing specialized compressors: tools such as JARVIS3 retain a clear advantage because they include explicit copy and repeat mechanisms. The value of genomic LMs is different. They provide reusable learned probability models that capture local composition, motif-like structure, coding-region regularities, and feature-associated predictability without hand-engineered genomic rules. A plausible practical path is therefore hybrid rather than competitive: explicit repeat/copy models for long duplicated segments, combined with LM-derived probabilities for the non-repeat residual and for biological interpretation.

Limitations. LLM compression is slow and not currently deployment-ready. Compressed sizes exclude model weights and tokenizer files, and decompression requires bit-exact access to the same weights, kernels, and quantization scheme. The human result is partly in-distribution because human sequence appears in the pretraining corpus. We evaluate only three genomes and fix the GPT-2-small architecture, so broader generalization and architecture-level effects remain open.

Future work. A controlled context-length ablation within DNAGPT2 would be the appropriate next experiment for isolating the role of context from architecture, tokenization, and training data. Hybrid compressors that combine LLM local distributions with explicit long-range repeat models are another natural next step. Online adaptation, as in NNCP, may improve ratios at higher cost. Biologically, correlating IC tracks with chromatin accessibility, eQTLs, and comparative genomics across more species would test whether these patterns generalize.

## 6. Conclusion

We show that a compact family of GPT-2-small models, trained on a multi-species DNA corpus and paired with arithmetic coding, achieves DNA compression competitive with strong specialized tools when compression is used as a likelihood-based evaluation metric. A controlled tokenizer sweep suggests that small BPE vocabularies may be preferable for genomic compression, while our model benchmark shows that published long-context genomic LMs do not automatically yield lower next-token entropy. These results should be read as model evaluation and biological interpretation, not as an attempt to replace mature specialized compressors. Beyond compression, the same models yield a genome-wide information-content map whose structure aligns with annotated biology.

### Software and Data

Code for training, compression, and analysis is available at https://github.com/ML-Bioinfo-CEITEC/llm_and_compression. The DNAGPT2 model collec-tion is available at https://huggingface.co/collections/vojtam/dnagpt2-68cda6809ef5f74179ca0e27. The interactive information-content map is available at https://genomeinfo.dyn.cloud.e-infra.cz/.

## Acknowledgements

We thank the anonymous reviewers for their constructive feedback. Computational resources were provided by the e-INFRA CZ project (ID:90254), supported by the Ministry of Education, Youth and Sports of the Czech Republic.

## A. Training Hyperparameters

**Table 6.** DNAGPT2 training hyperparameters.

| Group | Hyperparameter | Value |
| --- | --- | --- |
| Data | Total training tokens | $1.05 \times 10^{10}$ |
| Data | Epochs | 1 |
| Batch | Total batch size | $2^{19}$ tokens |
| Batch | Micro-batch size | 16 |
| Batch | Sequence length | 1024 |
| Batch | Gradient accumulation | 32 |
| Batch | Optimizer steps | 20,098 |
| Schedule | Max LR / min LR | $8 \times 10^{-4}$ / $8 \times 10^{-5}$ |
| Schedule | Warmup / decay | 1,000 / 19,098 steps |
| Optimizer | Algorithm | AdamW |
| Optimizer | Weight decay | 0.1 |
| Optimizer | Betas | (0.9, 0.95) |
| Precision | Mixed precision | bf16 |
| Hardware | GPU | 1 × NVIDIA A40 |
| Hardware | Wall-clock time | $\approx 7.5$ h |

## B. Additional Information-Content Views

## C. Supplementary Figures

**Figure 7.**
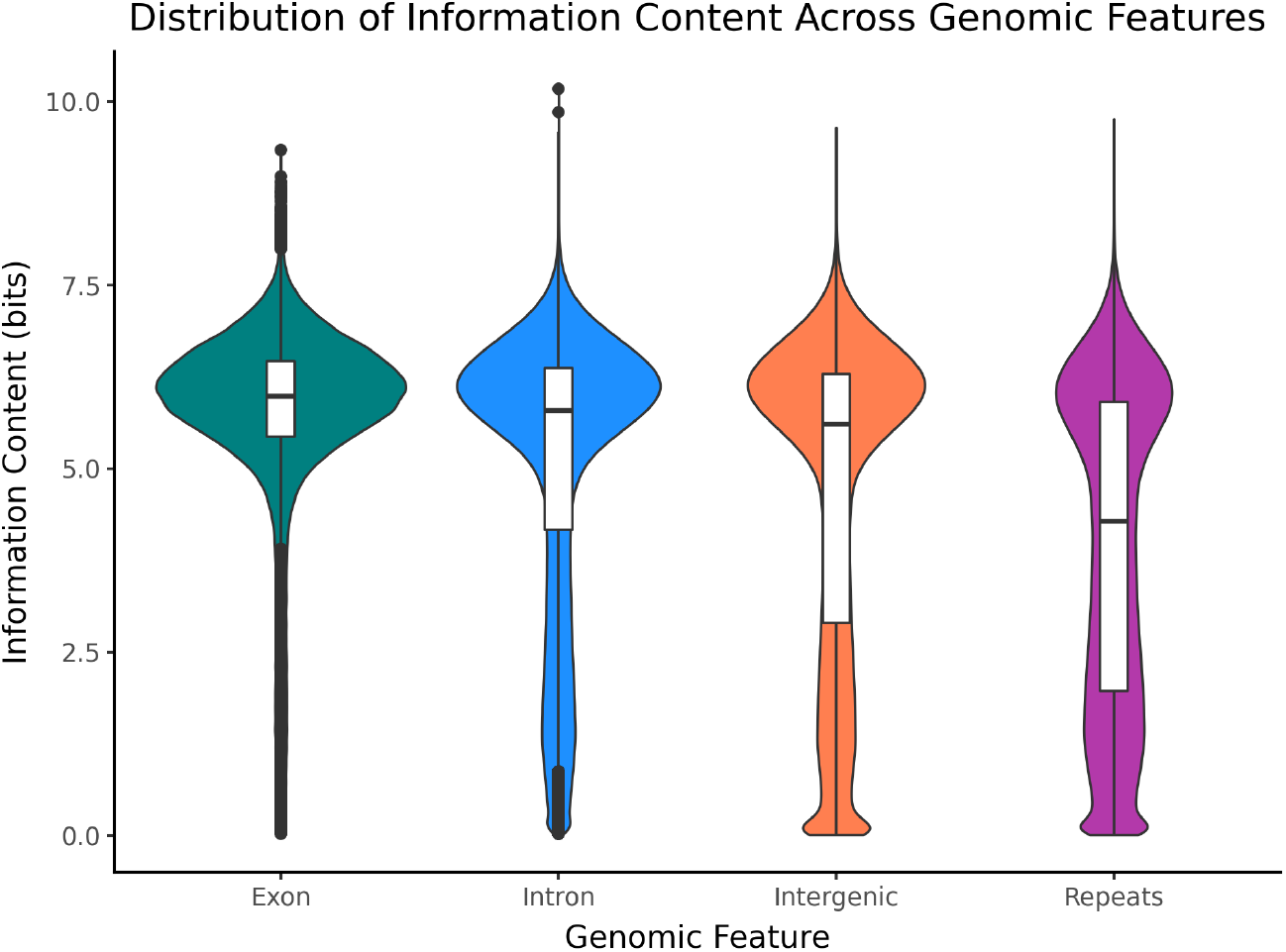
Variant of Figure 5 pooling all RepeatMasker repeat families and reporting raw negative log-probability per token (not normalized by token length).

**Figure 8.**
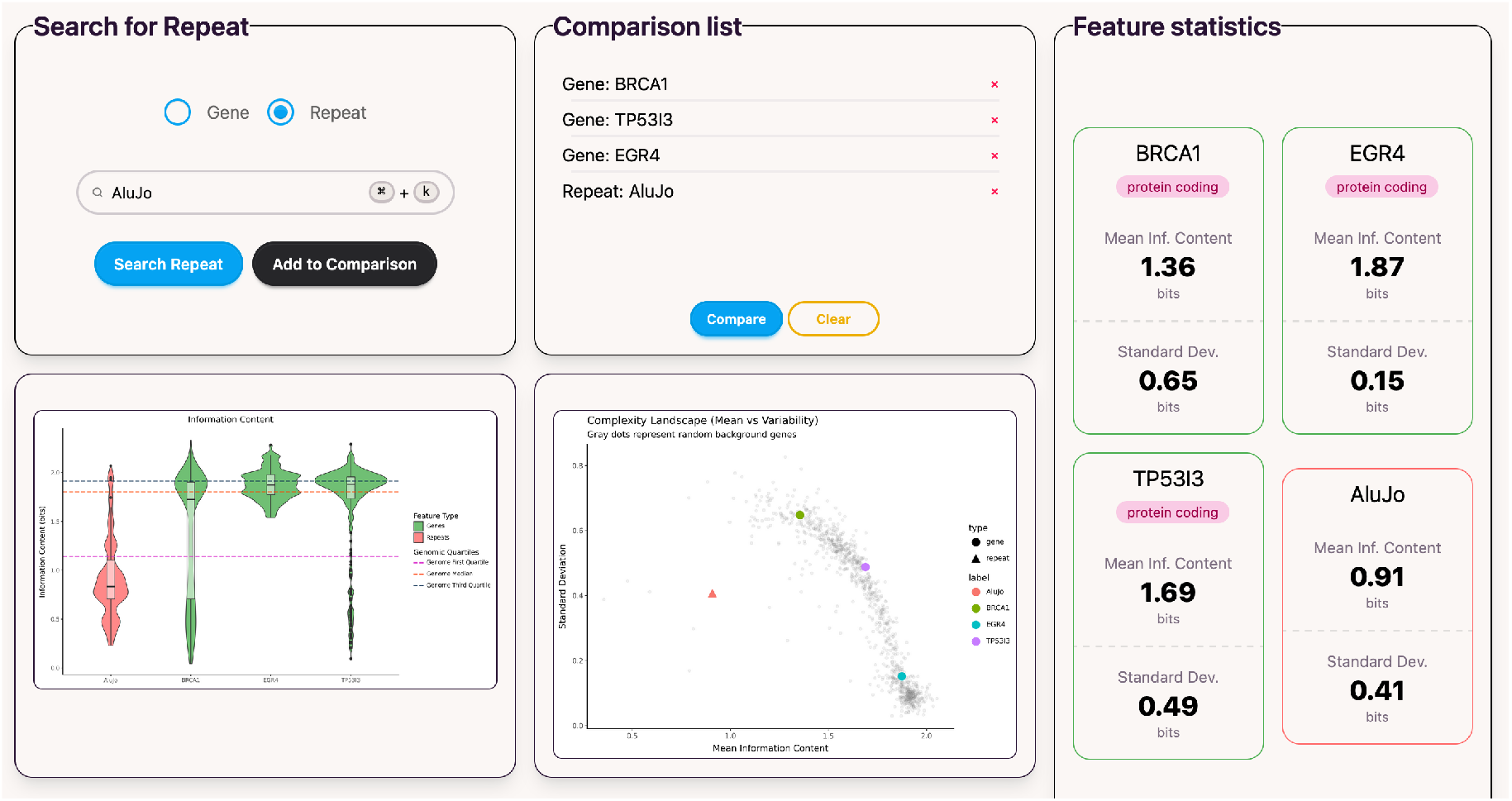
Interactive analysis interface for comparing user-selected genes and repeat families.

**Figure 9.**
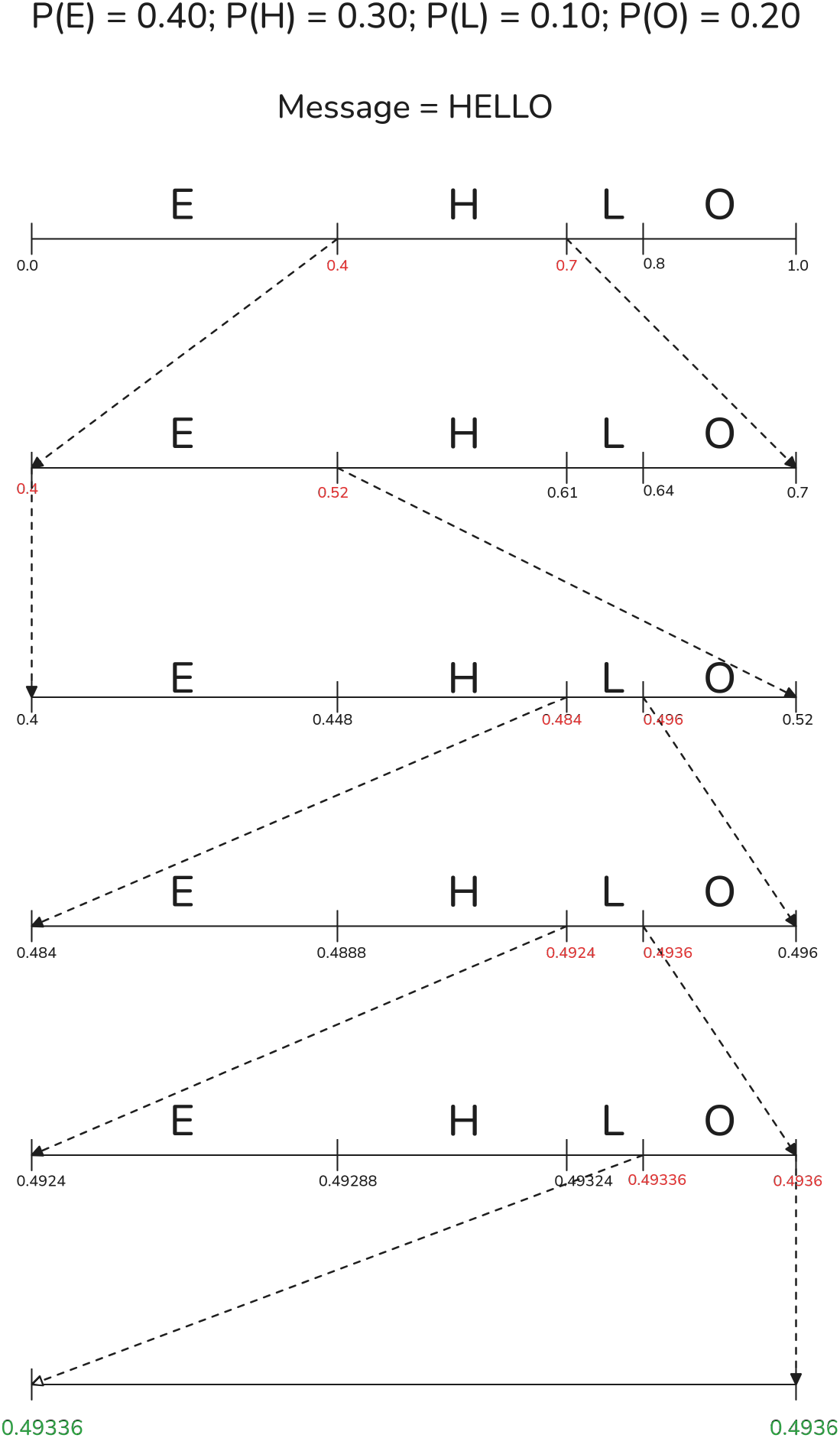
Arithmetic coding of “HELLO”. The unit interval is narrowed by each symbol; any binary fraction in the final interval decodes back to “HELLO”. Our pipeline performs the same interval narrowing using conditional distributions supplied by DNAGPT2.

## Footnotes

1 Abbreviated as M. llanfair… in tables and subsequent text.

1 Reported bpb measures the bitstream only. An 86M-parameter bf16 model is approximately 170 MB, which would add roughly 0.45 bpb of side-information overhead on a 3 Gb human genome if not amortized. We treat the model as side information throughout, consistent with using compression as a likelihood-based evaluation metric rather than a deployable storage system.

2 Times for non-LM rows are reported by Cobilab on their hardware; LLM rows were measured on a single NVIDIA A40 and are not strictly hardware-comparable.

